# Restrander: rapid orientation and QC of long-read cDNA data

**DOI:** 10.1101/2023.05.02.539165

**Authors:** Jakob Schuster, Matthew Ritchie, Quentin Gouil

**Author notes:** Correspondence |.

## Abstract

In transcriptomic analyses, it is helpful to keep track of the strand of the RNA molecules. However, the Oxford Nanopore long-read cDNA sequencing protocols generate reads that correspond to either the first or second-strand cDNA, therefore the strandedness of the initial transcript has to be inferred bioinformatically. Reverse transcription and PCR can also introduce artefacts which should be flagged in data pre-processing.

Here we introduce *Restrander*, a lightning-fast and highly accurate tool for restranding and quality checking long-read cDNA sequencing data. Thanks to its C++ implementation, *Restrander* was faster than Oxford Nanopore Technologies’ existing tool *Pychopper*, and correctly restranded more reads due to its strategy of searching for polyA/T tails in addition to primer sequences from the reverse transcription and template-switch steps. We found that restranding improved the process of visualising and exploring data, and increased the number of novel isoforms discovered by *bambu*, particularly in regions where sense and antisense transcripts co-occur.

The artefact detection implemented in *Restrander* quantifies reads which do not have the correct 5’ and 3’ ends, a feature which is useful in quality control for library preparation. *Restrander* is pre-configured for all major cDNA protocols, and can be customised with user-defined primers. *Restrander* is available at https://github.com/jakob-schuster/restrander

## Introduction

When performing gene expression analysis via RNAsequencing (RNA-seq), preserving information about the direction of the RNA transcript is important, especially in areas of overlapping transcription (1). With short-read sequencing, strand information is commonly preserved through selective degradation of the second strand cDNA, marked with dUTP (2). With Pacific Biosciences (PacBio) Iso-seq protocol, the polymerase read is a concatemer of alternating forward and reverse strand sequences, due to sequencing a circular template. Reads are oriented during the demultiplexing and adapter trimming step with *Lima*. With Oxford Nanopore Technologies (ONT), direct-RNA sequencing is inherently stranded since only the actual transcript is sequenced; by contrast, either the first or second-strand cDNAs may be sequenced in the direct-cDNA and PCR-cDNA protocols, resulting in both forward and reverse reads that have to be ‘restranded’ bioinformatically. Other protocols producing full-length double-stranded cDNA followed by nanopore sequencing (e.g. (3–5)) require the same restranding process. Library preparation for cDNA sequencing (Fig. 1A) adds different oligonucleotide sequences on the ends of each read. The template-switching oligo (TSO, also known as Strand-Switching Primer SSP) is found on the 5’ end of forward reads, and on the 3’ end of reverse reads (as a reverse complement). The sequence of the reverse transcription primer (RTP, also known as oligo(dT) VN primer or VNP in some protocols), is conversely found at the 3’ end of forward reads (as a reverse complement) and the 5’ end of reverse reads. Thus, when reads are sequenced from the 5’ end, forward reads and reverse reads can be differentiated by the order in which the TSO and RTP appear. In addition to these primers, polyA tails are present near the end of forward reads, while complementary polyT tails are found near the start of reverse reads. Searching for these tails as well as TSO/RTP sequences provides a more confident restranding process.

**Fig. 1.**
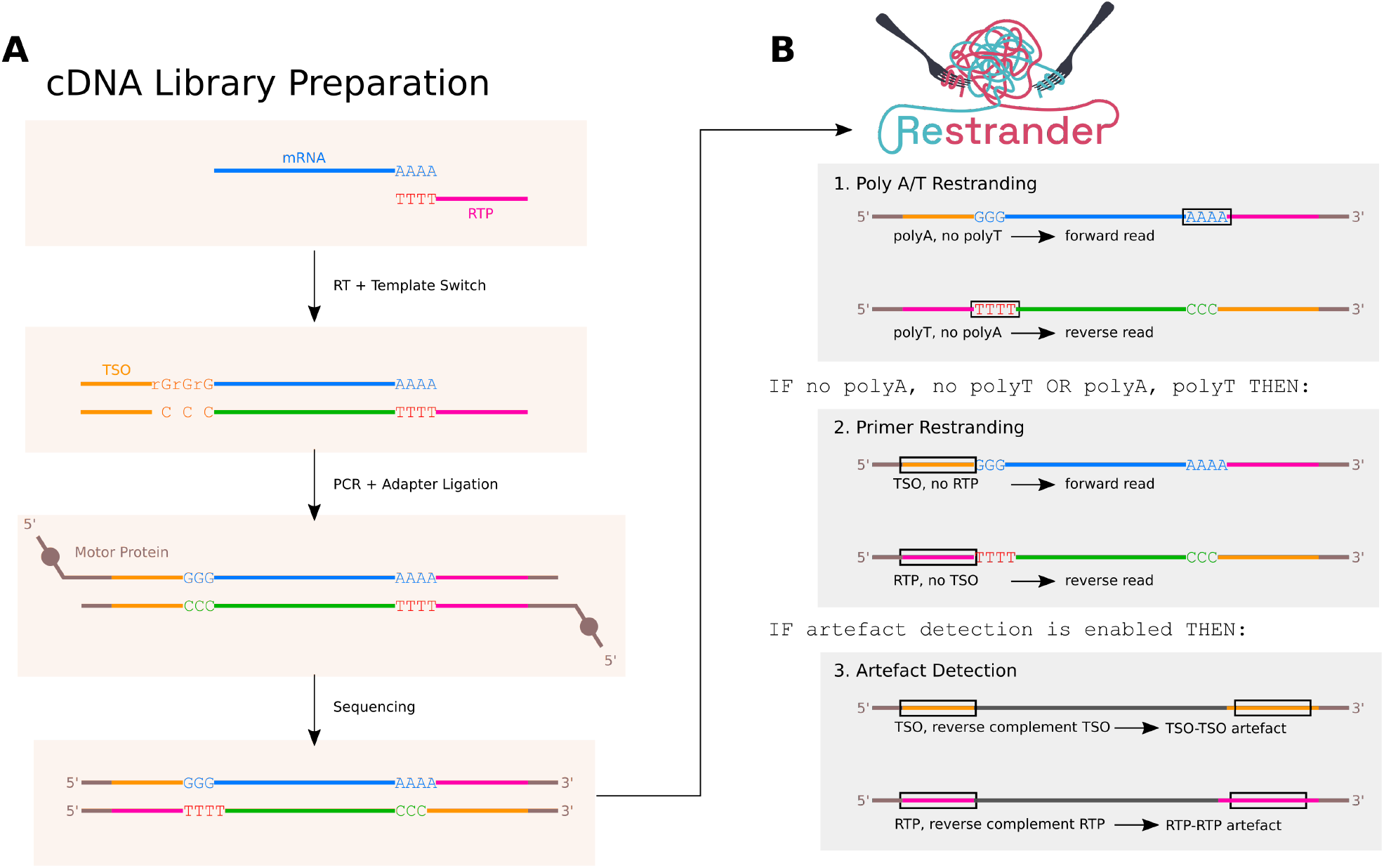
**(A)** Schematic of cDNA library preparation and sequencing. Forward reads are characterised by a lead TSO (template switch oligo) sequence, and a polyA-tail followed by the reverse complement of the reverse transcription primer (RTP). Reverse reads lead with RTP, a polyT stretch, and end with the reverse complement of the TSO. Different protocols (PCB109, PCB111, 10X Genomics, etc.) introduce variations but the principle remains the same. **(B)** The default processing pipeline used by *Restrander* to classify the direction of one read. If a read can be restranded using only the easily located polyA/T-tails, no further processing need take place. This optimisation considerably speeds up the process without significantly impacting number of incorrect classifications.

A number of tools exist for read orientation, including *ReorientExpress*, which uses a machine learning backend (6), and *primer-chop*, which employs fuzzy string search to locate primers (7). Third-party scripts have also been proposed (8). *Pychopper*, developed by ONT, is the most common tool for restranding reads from `fastq` files from direct-cDNA or PCR-cDNA nanopore runs, and has two backends one machine learning backend, and one fuzzy string search backend. Due to its widespread usage, *Pychopper* will be used for benchmarking in this paper.

While a variety of restranding tools are available, most have limited computational performance due to implementation in Python. This causes long processing times across large volumes of data. Additionally, for machine learning-based tools, adopting a new sequencing protocol requires the model to be retrained. For tools which employ fuzzy string search, accuracy in restranding reads could be improved by parsing for polyA/T tails in addition to primer sequences.

Here, we introduce *Restrander*, a more powerful and flexible restranding tool. We show that *Restrander* achieves greater accuracy, correctly restranding a higher proportion of reads while being one order of magnitude faster compared to *Pychopper*. Additionally, *Restrander* classifies artefactual reads to provide useful QC metrics and ensure only high-quality reads are taken for downstream processing. We propose *Restrander* as an integral step of nanopore transcriptomic pipelines, between basecalling and mapping.

## Materials and Methods

### A. Datasets

For all performance testing and analysis in this paper, we used a PCR-cDNA dataset of lung adenocarcinoma cell lines (9) which included synthetic RNA spike-in (sequins (10)), deposited under accession GEO GSE172421. Reads generated with SQK-PCS109 and PBK004 kits on PromethION R9.4.1 flow cells were basecalled using *guppy* v5.1.13+b292f4d13 with parameters `-c dna_r9.4.1_450bps_sup_prom.cfg --trim_strategy none --min_qscore 10 --barcode_kits SQK-PCB109 --do_read_splitting`. For the demonstration of accuracy on ground-truth data, we focused on a subset of 7M barcode01 pass reads, corresponding to cell line NCI-H1975 and sequins *mixA*. For speed testing, we concatenated reads from several barcodes to form `fastq` files of arbitrary size, producing files with 1M, 2M, 4M, 8M and 16M reads.

To test the compatibility of *Restrander* with common fulllength cDNA protocols, and to evaluate its artefact detection feature, we collected datasets from: rat single B cells prepared with the NEBNext Low input/single-cell kit for NAb-seq (4) (ENA Project PRJEB51442), single-cell longread RNA-seq from mouse muscle stem cells prepared with 10x Genomics Chromium 3’ kit (3) (GEO GSE154868) and PCR-cDNA libraries from human cell line THP-1 prepared with Oxford Nanopore Technologies SQK-PCB111 kit (ENA Project PRJEB60282).

### B. Alignment and transcript discovery

*minimap2* (2.17) was used to align reads to the reference genome concatenated with synthetic spike-in, with parameters `-ax splice`. *bambu* (1.0.3) (11) was run with default parameters.

### C. Performance comparison

*Restrander* (1.0.0) and *Pychopper* (2.7.3) were run on Intel(R) Xeon(R) CPUs E5-2690 v4 @ 2.60GHz (Broadwell), with 12 CPUs and 32GB of memory allocated per process. The processes were timed using wall clock time as measured by the Unix `time` command, with default parameters. Since *Pychopper* can run on multiple threads, it was given the default 8 threads. *Pychopper* has two separate computational backends. The default backend uses a machine learning model to classify reads, while the second uses the ‘edlib’ library to perform approximate string searching, employing similar methods to *Restrander* (12). Both backends were tested separately against *Restrander. Pychopper* was run with default parameters. *Restrander* was run with the default configuration for PCB109 data.

### D. Additional software

For generating the figures in this paper and performing basic analyses, R (4.2.1) (13) was used, in conjunction with the *ggplot2* (3.3.6) (14), *tidyr* (1.2.0) (15), *patchwork* (1.1.1) (16), *cowplot* (1.1.1) (17), *Gviz* (1.40.1) (18), *ggplotify* (0.1.0) (19), *GenomicAlignments* (1.32.0) (20), *rtracklayer* (1.56.1) (21) and *RColorBrewer* (1.1-3) (22) packages.

## Results

### E. *Restrander* Implementation

*Restrander* parses an input `fastq`, infers the strand of each read and prints to an output `fastq`. The strand of each read is recorded, and each read from the reverse strand is replaced with its reversecomplement, ensuring all reads in the output have the same orientation as the original transcripts.

In a typical cDNA-seq analysis pipeline, *Restrander* would be applied after basecalling, and before mapping. In the analysis for this paper, `fastqs` produced by *Guppy* were fed into *Restrander*, and then the restranded `fastqs` were used with *minimap2*. Only well-formed reads are included in the main output file; reads whose strand cannot be inferred are filtered out into an “unknown” `fastq`, to be handled separately by the user. If *Restrander* is configured to detect artefacts, these artefactual reads will also be placed in the “unknown” `fastq`.

The method used to process each read can be customised via the configuration file. Users can provide custom primer sequences and error rates, and modify parameters such as artefact detection and output verbosity to suit their needs. This enables *Restrander* to work across various protocols and contexts. Configuration files are provided for Oxford Nanopore Technologies SQK-DCS109, SQK-PCB109/PCS109, SQK-PCB111/PCS111; New England Biolabs NEBNext SingleCell/Low Input; 10x Genomics Chromium Single-Cell 5’ and 3’ kits. *Restrander* can also be run on adapter-trimmed reads, which have polyA/T tails without primers.

`./restrander input.fq.gz output.fq.gz config.json`

#### E.1. Restranding process

Using the default configuration, *Restrander* first searches for polyA/T tails (Fig. 1B). If the read strand cannot be conclusively inferred from this, then *Restrander* will search for TSO/RTPs. If the result is still inconclusive, the read is marked “unknown”. Although this initial polyA/T search speeds up restranding by about 5x overall, the user may specify to skip this step and always search for TSO/RTPs.

#### E.2. Implementation details

*Restrander* was written in C++ due to its excellent time performance over Python. The polyA/T tail classification algorithm searches for *n* consecutive A’s or T’s in the first *m* bases of the sequence. *n* = 10 and *m* = 200 by default, but both values are user-customisable. We found that if a polyA tail is not found within this hard limit *m*, it is very unlikely that one will occur at all. Thus, constraining our search to *m* does not negatively impact accuracy and ensures that extremely long sequences do not significantly slow down the restranding process. When polyA/T tail searching is unsuccessful, traditional approximate string matching will be performed to find TSOs and RTPs.

### F. Demonstration on ground-truth data

To demonstrate the accuracy of *Restrander*, we analysed a PCR-cDNA dataset of human cell lines spiked-in with sequins. Sequins are synthetic transcripts with known strandedness (10). By comparing the read classifications from *Restrander* and *Pychopper* with the reads’ alignments to the sequin reference transcriptome, we determined the accuracy of both *Restrander* and *Pychopper* on real data. Each set of inputs was repeated three times.

As shown in Fig. 2A, *Restrander* produced the most accurate results, correctly restranding 99.02% of reads, incorrectly restranding 0.67% of reads, and leaving 0.31% of reads marked unknown. *Pychopper*’s two backends were less accurate than *Pychopper*, with the machine learning and edlib backends correctly restranding 94.53% and 94.09% of reads respectively. Across a realistic dataset of 150M reads, even a difference of 5% would result in an additional 7.5M potentially useful reads.

**Fig. 2.**
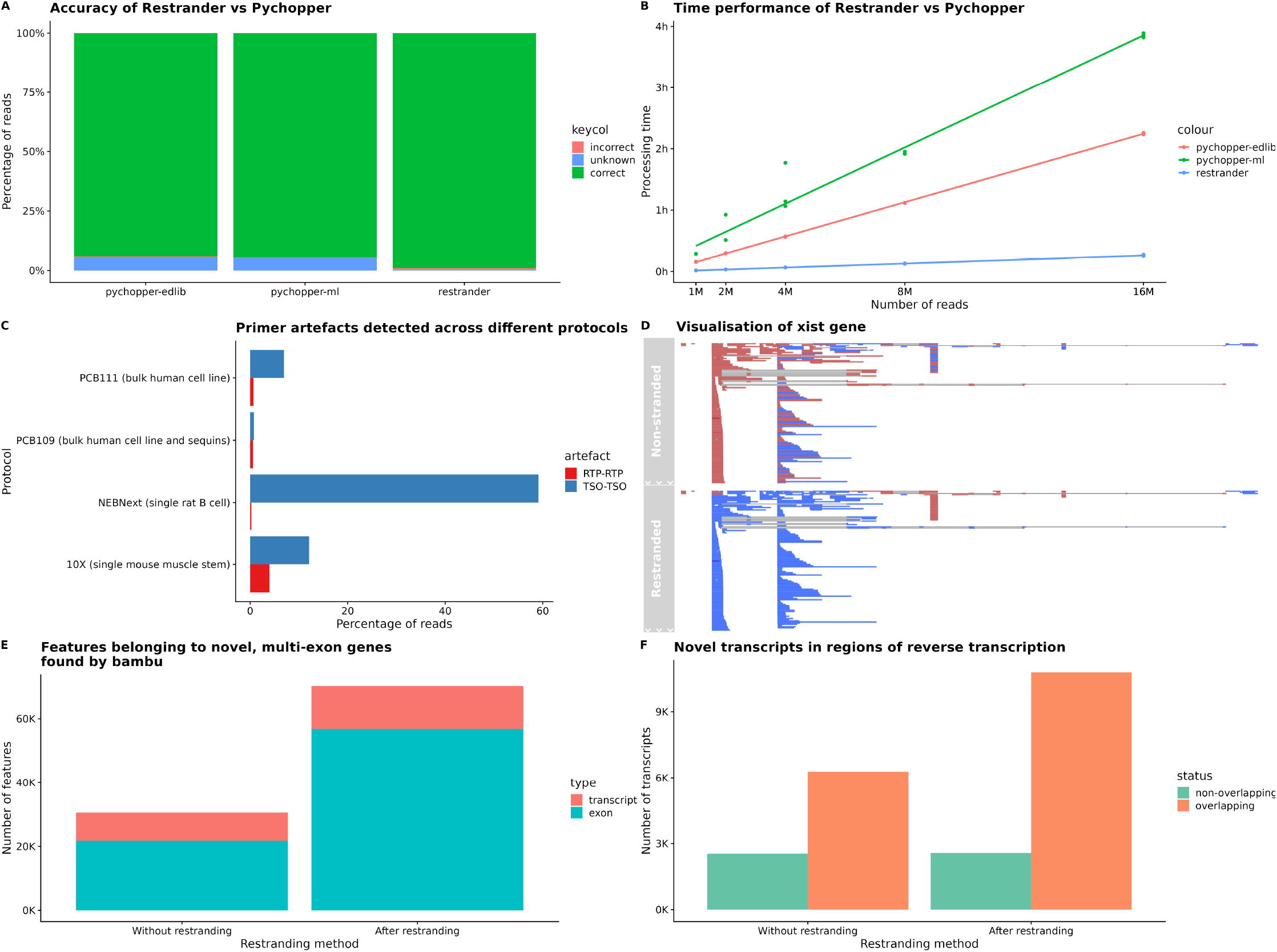
**(A)** Comparison of ground-truth accuracy of *Pychopper* and *Restrander*. **(B)** Comparison of time performance of *Pychopper* and *Restrander*. **(C)** Demonstration of artefact detection across various protocols and datasets. **(D)** Visualisation of a subsection of the *XIST* gene, with read directionality made clearer by the restranding process. **(E)** Number of features found by bambu on non-stranded vs restranded data, only considering features from novel, multi-exon genes. **(F)** Number of transcripts found by bambu originating from novel, multi-exon genes, comparing non-stranded and restranded data, and quantifying the number of transcripts which overlap with other transcripts on the opposite strand, as an indicator of sense and antisense transcript co-occurrence.

*Restrander*’s superior performance was likely due to the fact that, before searching for TSO and RTPs, it searches for polyA and polyT tails, taking advantage of a useful feature overlooked by both *Pychopper* backends.

Additionally, with default settings, *Restrander* categorises reads more confidently than *Pychopper*, leaving fewer unknown reads (*Restrander*’s 0.31% vs *Pychopper* ml’s 5.12% and edlib’s 5.57%) but resulting in more incorrect classifications (*Restrander*’s 0.67% vs *Pychopper* ml’s 0.35% and edlib’s 0.34%). If this is undesirable and the user would prefer to sacrifice volume of usable data for more certainty in read strandedness, they are recommended to run *Restrander* with a low error rate. This parameter and several others can be tweaked in the `json` configuration file.

### G. Speed comparison

*Restrander* runs much faster than *Pychopper*, a difference which becomes pronounced across higher volumes of data (Fig. 2B). Out of the two *Pychopper* backends, the machine learning backend is the slowest, taking 3 hours and 53 minutes to process 16 million reads. The edlib backend took only 2 hours and 15 minutes. *Restrander* completed the same task in 15 minutes. On different computer architectures, performance will vary, but the relative 9x speedup of *Restrander* over *Pychopper* is expected to be consistent across different hardware contexts. In a typical PromethION flow cell sequencing run of 150M reads, it is estimated to take *Pychopper*’s edlib backend roughly 21 hours to fully restrand the data, delaying downstream analysis. *Restrander* would take only 2 hours and 20 minutes, making restranding a much less costly process. Given the volume of data involved, this step is worth optimising. This improvement is primarily due to *Restrander*’s implementation in C++, as well as some algorithmic tweaks to minimize the processing involved.

### H. Artefact detection

*Restrander*’s rapid identification of TSO and RTP sequences lends itself to simultaneously investigating artefacts in library preparation. These include TSO-TSO artefacts (sometimes referred to as TSO artefacts) that may constitute up to 50% of reads from single-cell cDNA libraries (5), and RTP-RTP artefacts (referred to as RTP artefacts) by second-strand cDNA priming at internal polyT tracts in the transcript (23). These malformed reads are marked “unknown” and separated from the rest of the data.

On our PCB109 data, 0.77% of reads were TSO artefacts, and 0.58% were RTP artefacts. Some datasets contained significantly more artefacts (Fig. 2C). In particular, all other datasets contained more TSO artefacts than RTP artefacts; in the NEBNext data, 99.54% of the artefacts were TSO, affecting over 50% of all reads. As the datasets originate from a variety of RNA samples, we are not seeking to compare the prevalence of artefacts across different protocols, but instead demonstrate how *Restrander*’s artefact detection can be used to analyse primer-related artefacts that can be very prevalent.

### I. Data exploration

Restranding allows for clearer visualisation of RNA-seq data, particularly for regions where sense and antisense transcripts co-occur. To understand strange analysis results or inspect features, it is often useful to visualise data directly in IGV (24), or similar software. Restranded data can be more informative than non-stranded data for researchers to explore directly (Fig. 2D).

### J. Impact on isoform detection

To investigate the impact of restranded vs non-stranded data on downstream processing, we analysed a lung adenocarcinoma PCR-cDNA dataset aligned to the human genome reference CHM13 using *minimap2*. Isoform detection was performed using *bambu* in either unstranded or stranded mode. Single-exon genes found by *bambu* were discarded, as they were deemed low confidence. We hypothesised that stranded data would provide *bambu* with more informative reads and allow for more reliable isoform discovery, particularly in regions of the genome with sense and anti-sense transcripts co-occurring. Restranding allowed *bambu* to find a greater number of new features (exons and transcripts), particularly features from novel multi-exon genes; 70,234 features in stranded data versus 30,564 in non-stranded data (Fig. 2E). Of the transcripts from novel multi-exon genes, we observed that the restranded data had a greater number and higher proportion of transcripts from one strand overlapping with features found on the other (80.75% of new transcripts vs 71.12% for nonstranded data), suggesting that *bambu* found more features in areas of convergent transcription with the stranded data (Fig. 2F).

## Conclusions

In this paper, we present *Restrander*, a novel bioinformatic tool for quality control and transcript directionality inference of nanopore cDNA data. The software is open source and can be easily downloaded from GitHub under the MIT License. To assist users, a vignette is provided with a recommended workflow, making *Restrander* simple and easy-to-use.

*Restrander* is compatible with all common nanopore cDNA sequencing protocols, and additional configuration files can be used to accommodate new protocols and use cases. Our evaluation shows that *Restrander* outperforms *Pychopper* in both speed and accuracy in inferring transcript strand, which is critical for isoform annotation and quantification. *Restrander* offers an order of magnitude improvement in speed over *Pychopper*, and multi-threading could further enhance its performance.

Long-read sequencing technology has transformed transcriptomics, enabling the identification of full-length transcripts and alternative splicing events. However, the data generated can contain artefacts and be challenging to analyze. *Restrander* addresses this by identifying and filtering out artefacts that may affect over 50% of reads in some cases, enabling more accurate transcript quantification and splicing analysis. We also discovered that using restranded data in downstream isoform discovery via *bambu* led to the identification of more exons and transcripts, particularly in regions of convergent transcription. These novel features will require validation, which is beyond the scope of this manuscript.

We envision *Restrander* becoming a standard step in nanopore cDNA pipelines, used after basecalling and prior to mapping. By providing high-quality and stranded reads to downstream tools, *Restrander* will facilitate more accurate transcript quantification and splicing analysis.

## Data Availability

All supporting data is available through the following accessions: ENA Project PRJEB60282, and previously published ENA Project PRJEB51442(4), GEO GSE154868(3) and GEO GSE172421(9).

## Funding

QG and MER are supported by Australian National Health and Medical Research Council (NHMRC) Investigator Grants (GNT2007996 and GNT2017257, respectively). This work was supported by funding from the Chan Zuckerberg Initiative DAF, an advised fund of Silicon Valley Community Foundation (Grant No. 2019-002443 to MER).

## Conflict of interest statement

None declared.

## ACKNOWLEDGEMENTS

We thank Kathleen Zeglinski for designing the *Restrander* logo.

## Notes

https://github.com/jakob-schuster/restrander

### Competing Interest Statement

The authors have declared no competing interest.

### Summary of Updates

Acknowledged ReorientExpress and primer-chop in the introduction. Discussed benefits of Restrander in relation to other programs generally, not just Pychopper.

